# DECENT: Differential Expression with Capture Efficiency adjustmeNT for single-cell RNA-seq data

**DOI:** 10.1101/225177

**Authors:** Chengzhong Ye, Terence P Speed, Agus Salim

**Affiliations:** Bioinformatics Division, Walter and Eliza Hall Institute of Medical Research, Parville VIC 3052; Department of Mathematics and Statistics, La Trobe University, Bundoora VIC 3086; Baker Heart and Diabetes Institute, Melbourne, VIC 3004; Department of Mathematics and Statistics, The University of Melbourne, Parkville VIC 3010

**Keywords:** Differential expression, single-cell RNA-seq, dropout, imputation

## Abstract

Dropout is a common phenomenon in single-cell RNA-seq (scRNA-seq) data, and when left unaddressed affects the validity of the statistical analyses. Despite this, few current methods for differential expression (DE) analysis of scRNA-seq data explicitly model the dropout process. We develop DECENT, a DE method for scRNA-seq data that explicitly models the dropout process and performs statistical analyses on the inferred pre-dropout counts. We demonstrate using simulated and real datasets the superior performance of DECENT compared to existing methods. DECENT does not require spike-in data, but spike-ins can be used to improve performance when available. The method is implemented in a publicly-available R package.

## Introduction

Recent developments in sequencing technology have enabled high-throughput whole-transcriptome profiling at single-cell resolution. Single-cell RNA-seq (scRNA-seq) allows the quantification of gene expression of thousands of individual cells in a single experiment. It has already led to profound new discoveries that could not be have been made using data from bulk transcriptome sequencing, ranging from the identification of novel cell types to the study of global patterns of stochastic gene expression (Kolodziejczyk et al., 2015) (Wagner et al., 2016). However, there are still many statistical challenges in drawing inferences from scRNA-seq data. Due to the small amount of starting material and the imperfect capturing of RNA molecules in current scRNA-seq experiments, failures to detect expressed transcripts in single cells is still common. This gives rise to to the characteristic dropout phenomenon in scRNA-seq data, in which a gene shows zero or very low abundance in a fraction of cells in spite of moderate to high expression in others (Hashimshony et al., 2012) (Finak et al., 2015) (Ramskold et al., 2012). Also, the dropout rates can vary between cells and across genes (Brennecke et al., 2013), showing as a major source of unwanted variation in scRNA-seq data, with the first principal component of raw counts typically exhibiting high correlation with the proportions of zero counts (Risso et al., 2018). This unique feature of scRNA-seq will hinder downstream analyses if not properly modeled. Lots of effort has been made in order to alleviate this issue, including specialized normalization methods (Lun et al., 2016) (Bacher et al., 2017), clustering algorithms (Zeisel et al., 2015) (Wang et al., 2017) (Kiselev et al., 2017), and methods for differential expression analysis (Kharchenko et al., 2014) (Finak et al., 2015) (Jia et al., 2017).

One way to resolve this is through explicit modeling of the capturing process and hence separating the biological variation of interest from unwanted variation in the experimental procedures. For instance, imputation methods (Huang et al., 2018) (van Dijk et al., 2018) are designed to recover the pre-dropout expression matrix by modeling the process from RNA molecule to read count. However, a difficulty in modeling the molecule capturing and dropout events is that this process is usually mixed up with other sources of technical variation, such as amplification and sequencing biases (Wagner et al., 2016). The unique molecular identifier (UMI) barcoding approach has become increasingly popular in scRNA-seq experiments as an effective way to address this issue (Islam et al., 2014) (Svensson et al., 2017). Random barcodes are attached to cDNA molecules during reverse transcription. Each individual molecule from a particular gene in each cell is expected to have a distinct UMI (Islam et al., 2014). Therefore, after sequencing, by counting UMI barcodes instead of reads *per se*, the resulting UMI counts will be a faithful representation of the original cDNA counts, with amplification and sequencing bias largely avoided. But the UMI count will still show as zero if a RNA molecule failed to convert to cDNA, or was completely lost in amplification and sequencing. As a consequence, the main source of technical variation left in UMI counts is the loss of molecules during the experimental procedure, namely, dropouts. Hence, UMI count data provides us with an opportunity to model the molecule capturing process in depth. Also, given the distinct features of UMI-based data, it is necessary to build specific models in order to perform statistical tests reliably.

Currently scRNA-seq experiments mainly focus on cell-wise analyses such as clustering and trajectory inference for studying heterogeneity within cellular populations (Zeisel et al., 2015) (Trapnell et al., 2014) (Qiu et al., 2017). Nevertheless, differential gene expression (DE), as one of the most common gene-wise analyses, still plays an essential role in complementing these analyses. For example, it is used to identify cluster-specific markers for identifying the cell types. It is also used to derive disease-associated gene signatures (Sun et al., 2018) (Zhao et al., 2017) (Savas et al., 2018). However, DE methods originally designed for bulk RNA-seq tend to produce unreliable results due to failing to account for the extra variation in single-cell data (Jia et al., 2017) (Van den Berge et al., 2018). Driven by this, a few DE methods have been designed specifically for scRNA-seq data. All of them use some strategy to deal with the large variation and amount of zero observations. However, most of them do not distinguish biological from technical factors that are causing the phenomenon. For example, SCDE (Kharchenko et al., 2014) uses a mixture model to distinguish counts affected by dropout from the rest of the data. This model almost always assigns a probability of one that a zero count belongs to the dropout component, in essence assuming all observed zeroes to be technical. MAST (Finak et al., 2015) uses a two-part generalized linear model in which the dropout rates are adjusted by the inclusion of the observed fraction of non-zero counts as a term in their regression model. This still does not differentiate the dropouts from real biological zeros. Additionally, the effect of dropout events is likely to be non-linear, especially for genes with low to moderate expression (Bacher et al., 2017), and so the inclusion of simple linear term that represents capture rates in the regression model is unlikely to be optimal. ZINB-WaVE (Van den Berge et al., 2018) uses a zero-inflated model directly fitted to the observed data to derive observation weights for adjusting bulk DE methods. Only Jia *et al.* (Jia et al., 2017) proposed a DE method, TASC, that relies on external RNA spike-in data (Jiang et al., 2011) to fit a technical variation model in order to explicitly cater for dropouts, thus enabling separation of the biological variation for DE analysis. They showed improved performance of their method compared with methods that perform DE analysis directly using the observed data. Note that the methods mentioned so far are not specifically designed for UMI-count data. There are two existing methods that considers the unique features of UMI-based experiments: Monocle2 (Qiu et al., 2017) and NBID (Chen et al., 2018). They both fit negative binomial models directly to the observed UMI count without any explicit modeling of dropouts.

Here we propose a novel model for the DE analysis of UMI-based scRNA-seq data. Leveraging the features UMI-count data, we are able to model the molecule capturing process precisely. We build a dropout model to account for the gene- and cell-specific properties of molecule capturing. This allows us to perform DE analysis on the inferred pre-dropout distributions of RNA molecules. We named our method **D**ifferential **E**xpression with **C**apture **E**fficiency adjustme**NT** (DECENT). DECENT can use the external RNA spike-in data to calibrate the dropout model, but also works without spike-ins. In this paper, we describe the DECENT model and benchmark it against existing methods using both simulated data and four real UMI-based scRNA-seq datasets. The results showed improved performance of DECENT in various settings when compared to existing methods.

## Results

### Modeling extra-binomial variation in the capture process

ScRNA-seq data are noisy largely due to the complex experimental procedures. Each step introduces different sources of technical variation, which are further magnified by the low amount of starting material in a single cell. With the UMI barcoding approach, we are able to avoid part of the technical variation in the read counts caused by amplification and sequencing, primarily the over- and under-representation of RNA molecules (Islam et al., 2014) (Wagner et al., 2016). However, we still need to deal with the extensive loss of molecules that happen at all stages in a scRNA-seq experiment. For UMI-based experiments, we may simplify the procedures of the actual transcript counts in single cells turning into final UMI counts into a single capturing process, in which each RNA molecule is captured with a certain probability that we term capture efficiency. We aim to design a dropout model to describe this capture process, which will enable us to infer the unobserved pre-dropout RNA molecule counts and perform DE analysis on them. It is natural to consider capture efficiencies to be cell-specific, as molecules from the same cell are in the same reaction chamber (well or droplet) during the capturing process. Previous models have considered between-cell variation of capture efficiency as a major source of technical variation in scRNA-seq data (Grun et al., 2014). By further assuming the capture of each molecule is independent within a cell, we obtain the simplest dropout model, a binomial thinning process ***B***(*y*_*i* *j*_, *η*_*j*_), where *y*_*i* *j*_ is the unobserved pre-dropout molecule count of gene *i* in cell *j* and *η*_*j*_ is the molecule capture efficiency in cell *j*. We denote the observed UMI counts by *z*_*i* *j*_.

We now examine the plausibility of this simple dropout model using ERCC spike-in data. ERCC spike-ins are synthetic RNA molecules added to the initial RNA pool in transcriptome profiling assays to measure technical variation (Jiang et al., 2011). The nominal molecule count of each spike-in added per cell (*c*_*i*_) is known, with no biological variation between cells expected. We thus use a Poisson distribution with rate *c*_*i*_ to model pre-dropout spike-in molecule count *y*_*i* *j*_, where *c*_*i*_ is the nominal molecule count of spike-in *i*. This is to model the sampling noise due to dilution. The known pre-dropout distribution of spike-in data enables us to focus on examining the dropout model. In total, we investigated six ERCC spike-in datasets, including three plate-based (Tung et al., 2017) (Zeisel et al., 2015) (Grun et al., 2014) and three droplet-based experiments (Macosko et al., 2015), (Klein et al., 2015), (Zheng et al., 2017) (Supplementary Table 1). Taking the Tung *et al.* data as an example, we first estimated the capture efficiencies *η*_*j*_ under the hypothesized model. We then used deviance statistics to evaluate goodness-of-fit of this cell-specific binomial dropout model. It can be easily calculated as a log-likelihood ratio (See Supplementary Methods). Under the null hypothesis that the binomial dropout model is adequate, the deviances approximately follow a *χ*^2^ distribution of with degree of freedom (#spike-ins - #parameters). However, as shown in Fig.1a, the observed distribution of cell-wise deviances is evidently shifted towards the higher values, indicating a poor fit to the data.

**Figure 1:**
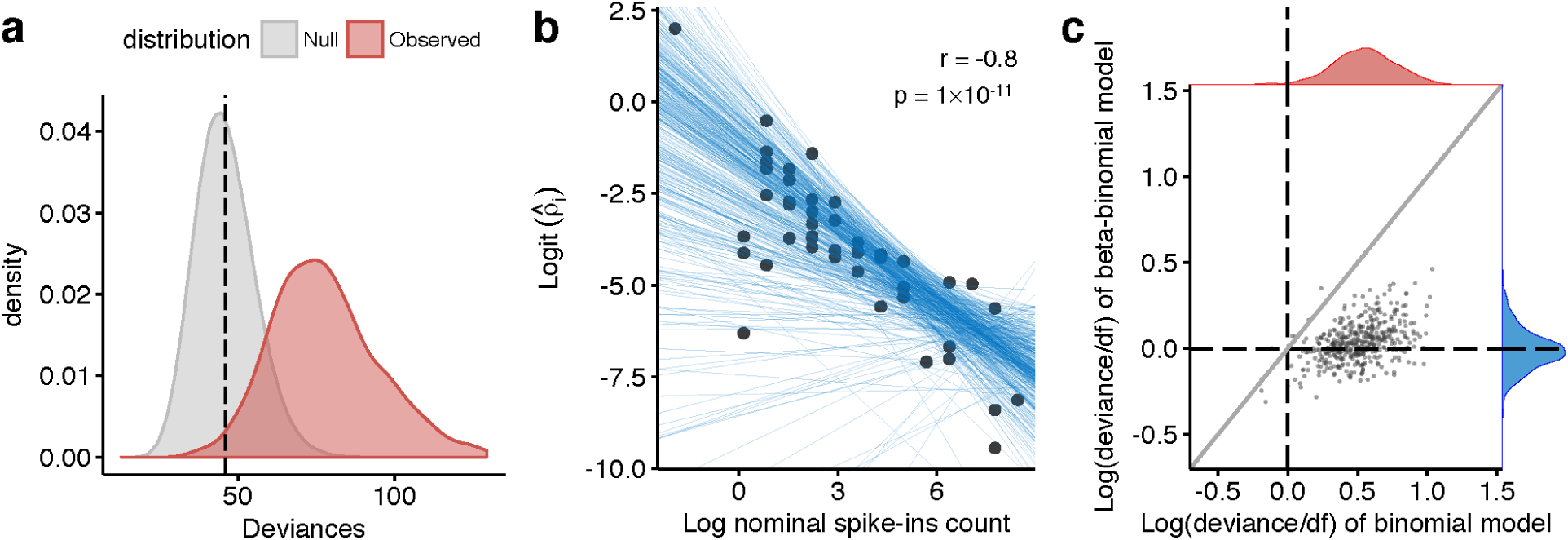
Modeling extra-binomial variation in the molecule capturing process. We evaluate the binomial and beta-binomial dropout models using the ERCC spike-in data from the Tung *et al.* experiment. (**a**) The observed distribution (red) of deviances with cell-wise binomial dropout model shows notable deviation from the expected χ^2^ distribution the under null hypothesis. This indicates inadequacy of the binomial dropout model. (**b**) Modeling the relationship between the spike-in nominal count *c*_*i*_ and the dispersion parameter ρ in the beta-binomial dropout model. If the parameter is estimated in a spike-in specific manner, a high correlation between the ρ_*i*_ estimates and the true pre-dropout mean abundance, namely the nominal count *c*_*i*_, can be observed, which are shown as black points. We build a cell-wise linear model to characterize this relationship. Each blue line represents a fitted cell-wise model, which is shown to adequately describe this relationship. (**c**): A scatter plot comparing the cell-wise deviances under the binomial and beta-binomial dropout models to assess goodness-of-fit. Deviances were standardized by dividing by the degrees of freedom to enable comparison, and logged. The blue and red marginal densities represent the observed distributions of deviances under the two models respectively. It can be seen that the beta-binomial dropout model fits better than the binomial model in the majority of the cells.

This shows the inadequacy of the cell-specific binomial dropout model. It could be due to the spike-in-wise variation of capture efficiency or a clumping of molecules in which the capture events are actually not independent within a cell. Further analyses suggests the former as a more probable main cause (Supplementary Fig.1). We model this extra variation by allowing capture efficiencies *η*_*j*_ to have a beta distribution with dispersion parameter *ρ* instead of being constant in a cell, resulting in a beta-binomial dropout model, *BB*(*y*_*i* *j*_, *η*_*j*_, *ρ*). Note that this is not the conventional parametrization of beta-binomial (See Methods). We then investigated whether a constant, cell-specific *ρ* is adequate. Towards this end, we looked at the variation of *ρ* across spike-ins. If we estimate *ρ*_*i*_ for each spike-in separately, a negative correlation between the spike-in-specific *ρ*_*i*_ estimates and the spike-in nominal count *c*_*i*_ can be observed (Fig.1b). This indicates a standard cell-wise beta-binomial dropout model with a single *ρ*_*j*_ for the all genes will not adequately describe the variation in the capture process. To deal with this, we constructed a simple logistic linear model for cell and gene-specific *ρ*_*i* *j*_:

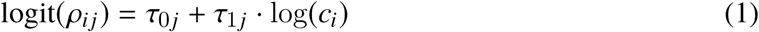

where **τ _*j*_** = (τ_0*j*_, τ_1*j*_) are the intercept and slope determining how *ρ*_*i* *j*_ depends on *c*_*i*_ in cell *j*. Using spike-ins, the estimation of τ**_j_** by maximum-likelihood is straightforward (see Methods). As a final examination, we fit both the binomial and this beta-binomial dropout models to the data and compared their goodness-of fit. Since the two models have different degrees of freedom, to facilitate comparison, we standardized and transformed the deviances of both models by forming log(*deviance*/*df*), which is expected to have a 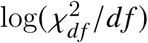 distribution with mean 0. As expected, we found cell-wise log(*deviance*/*df*) values under our beta-binomial model to be substantially smaller than those under the binomial dropout model for most of the cells (Fig.1c). We also observed that the distribution of *log*(*deviance*/*df*) values under our beta-binomial model are centered around 0, suggesting a good match to the null distribution. This indicates that our beta-binomial dropout model should satisfactorily account for variation in the molecule capturing process. The same analyses were carried out using the spike-in data from the other five experiments and similar results were obtained (Supplementary Fig.2).

### Inferring the distribution of pre-dropout molecule counts

We now start considering the model for RNA molecules from endogenous genes. We expect the external spike-ins and endogenous transcripts to have similar but not identical capture processes. Therefore, we use the same dropout model for endogenous genes with partially re-estimated parameters (See Methods for detail). To infer the pre-dropout molecule counts, we need to specify a distribution that characterizes them. We used a Poisson distribution to model the pre-dropout molecule count of spike-ins where no biological variation is expected. This is unlikely to be appropriate for endogenous genes. Instead, we chose to use the zero-inflated negative binomial (ZINB) distribution, which has been used in some scRNA-seq methods to model the observed counts (Risso et al., 2018) (Van den Berge et al., 2018). The ZINB distribution is a mixture of two components, a negative binomial distribution component and a structural zero component. The negative binomial (NB) distribution alone has been previously used in bulk DE methods (Robinson et al., 2010) (McCarthy et al., 2012) (Love et al., 2014) to model overdispersion in data due to gene-specific biological variation. In addition, we expect biological zeros at the single-cell level to be more abundant due to phenomena such as stochastic gene expression, state-dependent expression and heterogeneous cell composition, which are not observable in bulk RNA-seq experiments (Raj et al., 2006) (Shalek et al., 2013) (Buettner et al., 2015). The structural zero component is used to model these inflated biological zeros. These result in the DECENT framework that describes the molecule capture process. The pre-dropout RNA molecule count *y*_*i* *j*_ from gene *i* in cell *j* is assumed to follow a ZINB distribution with gene-specific dispersion and zero-inflation parameters. After molecule capturing, we observe an UMI count *z*_*i* *j*_ that is generated according to the beta-binomial dropout model (See Methods for detail). We use the DECENT model to distinguish biological from technical variation due to dropouts and perform differential expression analysis on the inferred pre-dropout distribution.

Next we investigated how well we can infer the pre-dropout counts by simulation studies. We simulated a dataset of 500 cells, 3000 endogenous genes and 50 detected spike-ins, with parameters empirically estimated from the Tung *et al.* dataset. We fit the DECENT model to this simulated data and then looked into two main features of the pre-dropout counts: proportion of zeros and variance. We calculated the gene-wise zero fractions and variances of the inferred pre-dropout counts and found the values are very close to those calculated using the actual pre-dropout counts (Fig.2a, b). We further calculated the expected pre-dropout count of each gene in each cell based on the fitted model and the observed data. Again, we found it to be highly consistent with the true count (Fig.2c).

**Figure 2:**
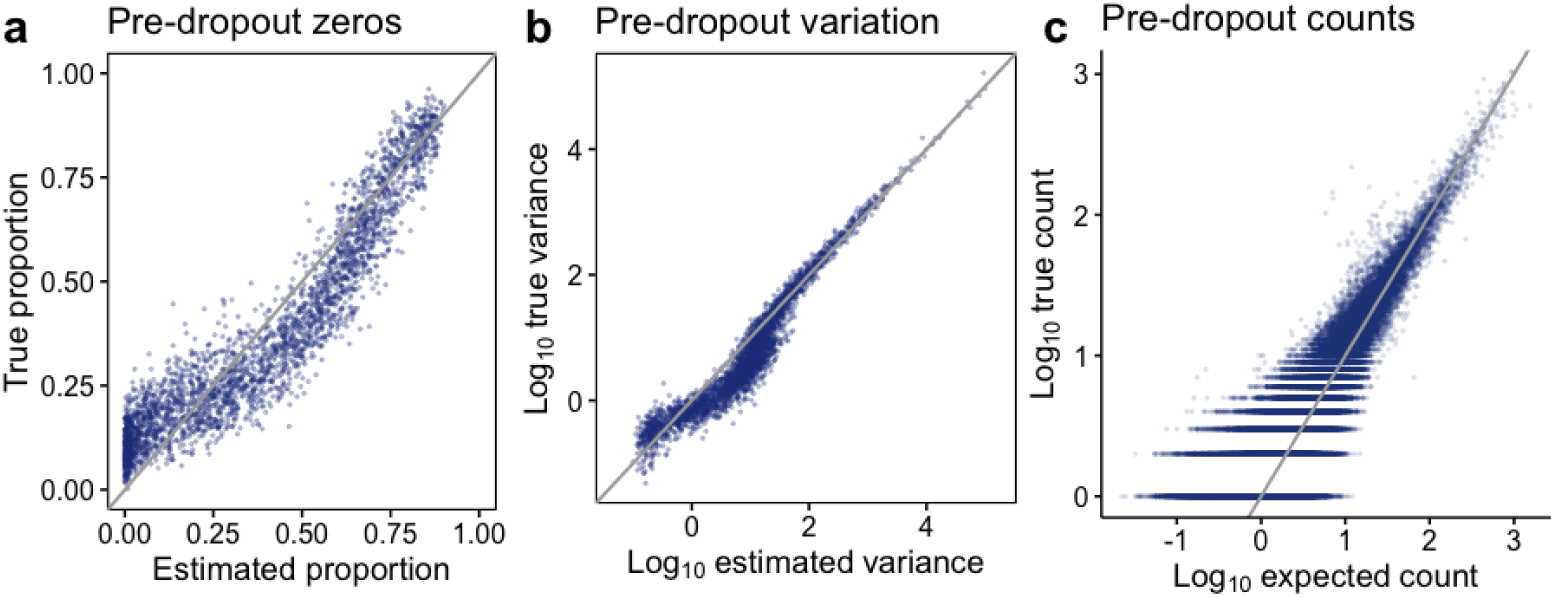
Inferring pre-dropout molecule counts in simulation. (**a**) Scatter plot comparing for each gene the estimated proportion of zeros of the fitted pre-dropout distribution with the true proportion of zeros in the pre-dropout counts. (**b**) Scatter plot comparing the expected variance of the fitted pre-dropout distribution with the true gene-wise variance in the pre-dropout counts. (**c**) Scatter plot comparing the expected value of pre-dropout count (see Supplementary methods for details) under the fitted model with the true pre-dropout counts. We showed a random subsample of 5 percent of all the non-zero counts. The estimated pre-dropout counts used to calculate (**a**) and were based on single imputation, i.e., drawing a single value from the conditional pre-dropout distribution for each gene and each cell given the parameter estimates and the observed data. The estimated pre-dropout counts shown in (**c**) were calculated as the expected value of the conditional pre-dropout distribution (See Supplementary Methods).

To examine whether there is overdispersion and zero-inflation in pre-dropout counts in reality, we used two scRNA-seq datasets where spike-ins are available (Zeisel et al., 2015), (Tung et al., 2017). Therefore, capture efficiencies could be estimated using the spike-ins to obtain reliable dropout models. To look for overdispersion, we first fit the DECENT model assuming an NB pre-dropout distribution to the data without considering zero-inflation. We found that without gene-specific dispersion parameters, the expected variances of most genes were noticeably lower than the observed values for the Zeisel *et al.* dataset. The extra variation was modeled by having the dispersion parameter (Supplementary Fig.3a). For the Tung *et al.* data, the expected variances without overdispersion parameters for most genes were already close to the observed values, showing little need for the extra parameter (Supplementary Fig.3b). This suggests overdispersion in pre-dropout counts is dataset-specific and depends on the amount of biological variability in the sample. The Tung *et al.* data used here are from one iPSC cell line where cells were highly homogeneous and hence lack biological variation. On the other hand, the Zeisel *et al.* data are from mouse brain tissue, which has a complex cellular composition. To check for zero-inflation, we then fitted DECENT models to the data assuming ZINB pre-dropout distributions. We performed chi-square goodness-of-fit test on both DECENT models with ZINB and NB to assess their adequacy. Consistent with previous findings (Vieth et al., 2017) (Chen et al., 2018), the majority of genes do not need to have a zero-inflated model. However we still found a small number of genes in both datasets in which models with ZINB provide a more adequate fit than NB (Supplementary Fig.4). Overall, the ZINB distribution provides a comprehensive solution for modeling the pre-dropout molecule counts. Zero-inflation and overdispersion in gene-wise pre-dropout counts turned out to be dataset-specific and gene-specific. In the cases where these effects are less prominent, the ZINB distribution will have low pre-dropout zero-inflation and/or dispersion parameter estimates, and effectively turn into NB, zero-inflated Poisson or Poisson distributions.

Additionally, we investigated a single-molecule fluorescence *in situ* hybridization (smFISH) dataset. The smFISH technology allows precise quantification of RNA molecules from a list of targeted genes. This technology can achieve near 100% sensitivity detection of the RNA molecules (Raj et al., 2008). In other words, the smFISH count data may be a good approximation to the pre-dropout molecule counts and should follow our assumed distribution. We used the data from an experiment that profiled 33 marker genes in mouse somatosensory cortex (Codeluppi et al., 2018). We examined three of the clusters identified by the authors, Oligodendrocyte Mature, Pyramidal L4 and Inhibitory Vip, finding most of the gene count distribution to be significantly overdispersed relative to the Poisson (Supplementary Fig.5a). Yet we did not find zero-inflated genes in these clusters. This is quite possibly because the targeted genes are all canonical markers, which are expected to mostly exhibit constitutive expression and hence unlikely to have inflated zeros caused by transcriptional bursting. However, heterogeneity within a population can also result in zero-inflation, which is common in actual DE analysis. We thus increased the heterogeneity within the groups by focusing on three major cell types, Oligodendrocytes, Pyramidal neurons and Inhibitory neurons. We then identified two, one and two out of the 33 genes to have significant zero-inflation (Supplementary Fig.5b).

### Benchmarking using simulated data

We next performed DE analysis using the simulated data and benchmarked DECENT against several existing methods. These includes SCDE (Kharchenko et al., 2014), MAST (Finak et al., 2015), Monocle2 (Trapnell et al., 2014) (Qiu et al., 2017), ZINB-WaVE adjusted edgeR (Van den Berge et al., 2018), and the standard edgeR (McCarthy et al., 2012) to represent bulk DE methods. We set a fraction of genes in the simulated data to have higher log fold-changes and used those as the reference genuine DEGs for benchmarking. Firstly, to assess the general ability of each method to distinguish between DEGs and non DEGs, we used the receiver operating characteristic (ROC) curves based on the nominal p-values produced by different methods. In actual DE analysis, usually only the low p-value region is of interest, so we used the partial ROC (pROC) curve (McClish, 1989) (Robin et al., 2011) focusing on the region with false positive rate smaller than 0.1. As shown in Fig.3a, DECENT outperformed all other methods in the simulation study. To further evaluate the level of false positives among the top discovered DEGs, we used the false discovery rate (FDR) curve, describing the fraction of false discoveries among the top n declared DEGs by each method. Again DECENT showed the smallest fraction of false discoveries consistently (Fig.3b).

**Figure 3:**
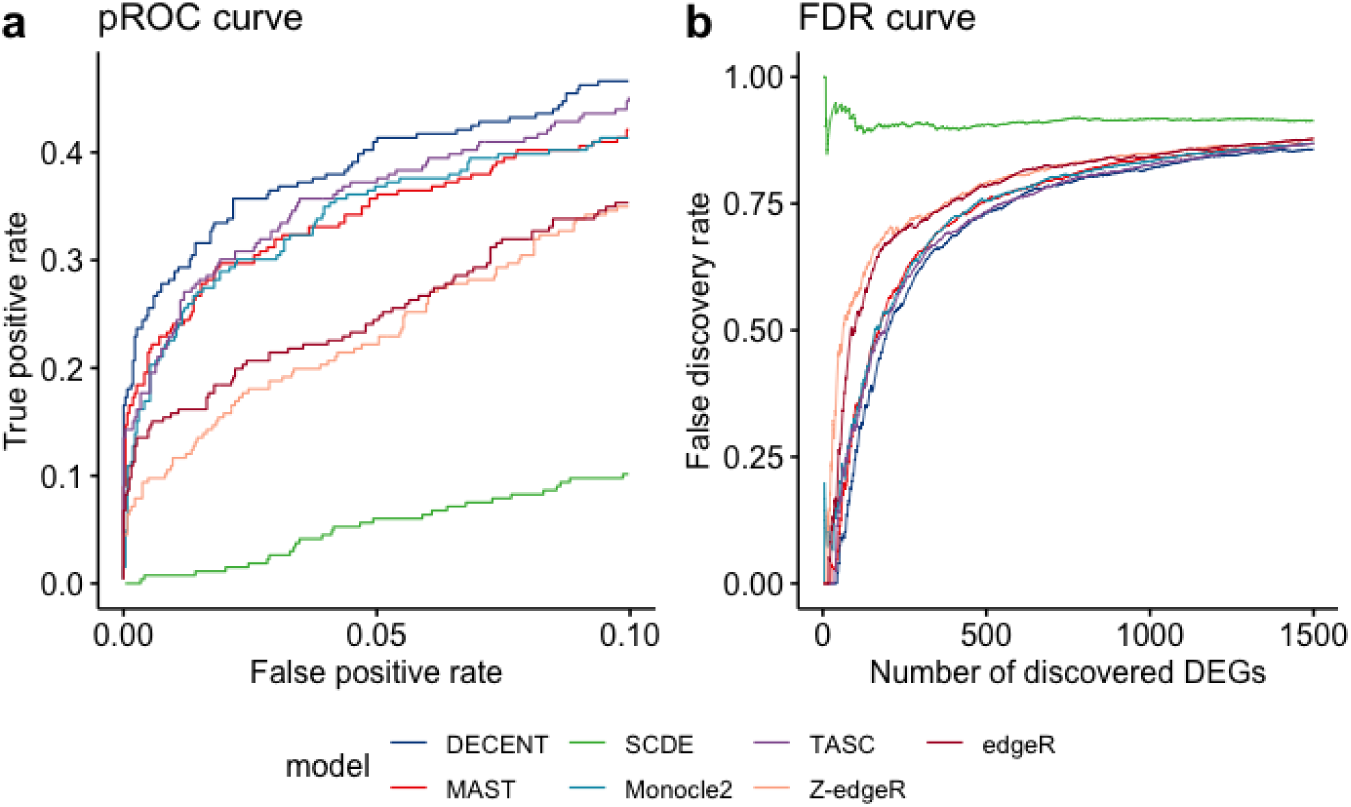
Differential expression analysis of simulated data. (**a**) Partial receiver operating characteristic curve for differential expression methods on the simulated data. (**b**) False discovery rate curves for differential expression methods on the simulated data. Both curves only focus on the low p-value region, since other regions were of little interest in actual DE analysis. Z-edgeR stands for ZINB-WaVE-adjusted edgeR.

Many existing scRNA-seq data datasets do not have spike-ins. Also, the recently popularized droplet-based technologies are incompatible with spike-ins. We therefore want to enable the usage of DECENT without spike-ins. To this end, we have developed a strategy to obtain functional dropout models only using information of endogenous genes. Basically, we assign ranked random capture efficiencies to each cells according the empirical distribution of the observed library size. We then fit the DECENT model assuming no spike-in information available. According to certain properties of our model, other components will compensate for the inaccuracy of capture efficiency estimates (Supplementary Fig.6, see Methods for details). To examine how this affect DE analysis, we set the range of the ranked random capture efficiencies to be either the same as (1x), half (0.5x) or one and a half (1.5x) the true range. We found DE analysis is mostly unaffected by using this strategy to obtain dropout models and robust to inaccurate capture efficiencies. Except the performance decreases slightly when the capture efficiencies are specified too high (Supplementary Fig.7). This is possibly because the unaccounted variation is so large that it goes beyond the extent to which the model can adjust itself. Therefore, we generally recommend setting smaller ranges of capture efficiencies.

### Benchmarking using real data

The simulation study has demonstrated the feasibility of our model mathematically. However, it cannot prove that our model assumptions or DE strategy are appropriate for genuine biological data and questions. Hence, we further benchmarked our model against existing methods using real datasets. The difficulty in benchmarking using real datasets is that the genuine DEGs are usually unknown. In order to obtain a credible list of genuine DEGs, we searched for scRNA-seq datasets that have matching bulk RNA-seq experiments, which means a bulk RNA-seq was also performed using cells from exactly the same tissues or cell lines. We found four such experiments in total that also used UMI. Then a DEG list derived from these bulk data can be used as the reference set for benchmarking. These includes two plate-based experiments and two droplet-based experiments, with different scales, sources of tissues or cell lines and observed proportion of zeros (Supplementary Table 2) (Tung et al., 2017) (Soumillon et al., 2014) (Savas et al., 2018) (Chen et al., 2018).

We again evaluated the performance using pROC and FDR curves. The same methods as the last section were benchmarked using all four datasets, except that we also applied TASC to the Tung *et al.* data where spike-ins are available. As shown in Fig.4 and 5, DECENT showed superior performance on all four datasets. MAST showed stable and generally acceptable performance across datasets, while the performance of SCDE appeared to be dataset-specific, showing inadequacy for droplet-based experiments. The Monocle negative binomial-based model based on observed UMI count did not show satisfactory performance. The ZINB-WaVE adjustment of edgeR did not show noticeable improvements over standard edgeR for three out of four datasets. But it remarkably outperformed edgeR on the Chen *et al.* data, where both molecule counts and the cell numbers were high. To demonstrate the merit of performing DE analysis using a inferred pre-dropout rather than the observed expression, we selected a few genuine DEGs in the Tung *et al.* data that are detected by our method and compared their expression levels between the two cellular groups using either the observed counts or inferred pre-dropout counts. We discovered that the differential expression between two groups became more prominent in the pre-dropout counts (Supplementary Fig.8).

**Figure 4:**
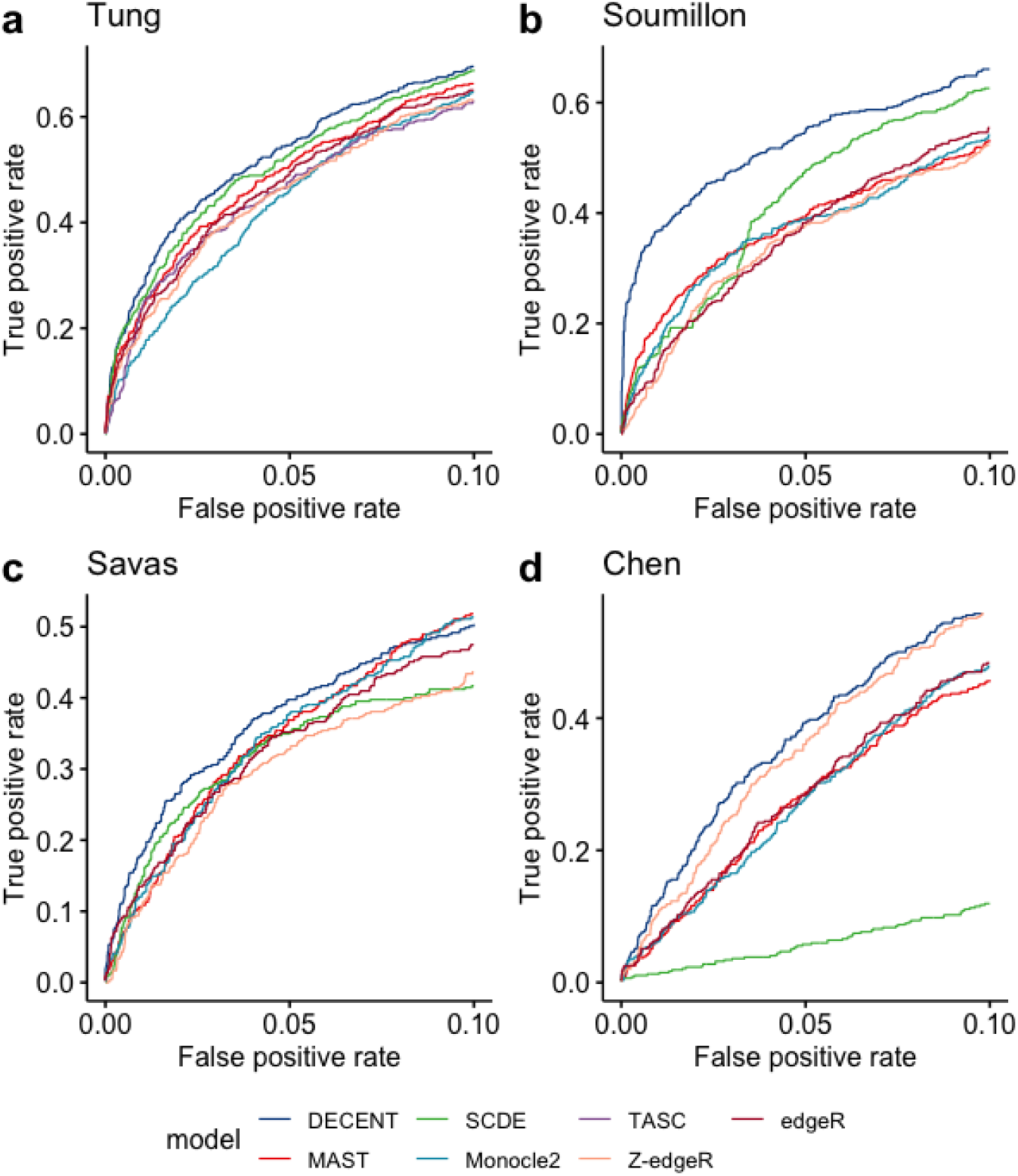
Partial receiver operating characteristic curves for differential expression methods on real datasets. Evaluating the performance of different methods by partial receiver operating characteristic curves using (**a**) Tung et *al.*, (**b**) Soumillon *et al.*, (**c**) Savas *et al.* and (**d**) Chen *et al.* datasets. DEGs from matching bulk RNA-seq data were used as gold-standard for benchmarking. DECENT achieves highest accuracy of identifying genuine DEGs in all four datasets. We used pROC to focus on the low p-value region with high specificity. DE methods are denoted by different colors. Z-edgeR stands for ZINB-WaVE-adjusted edgeR. TASC requires spike-ins and was only evaluated using the Tung *et al.* data.

**Figure 5:**
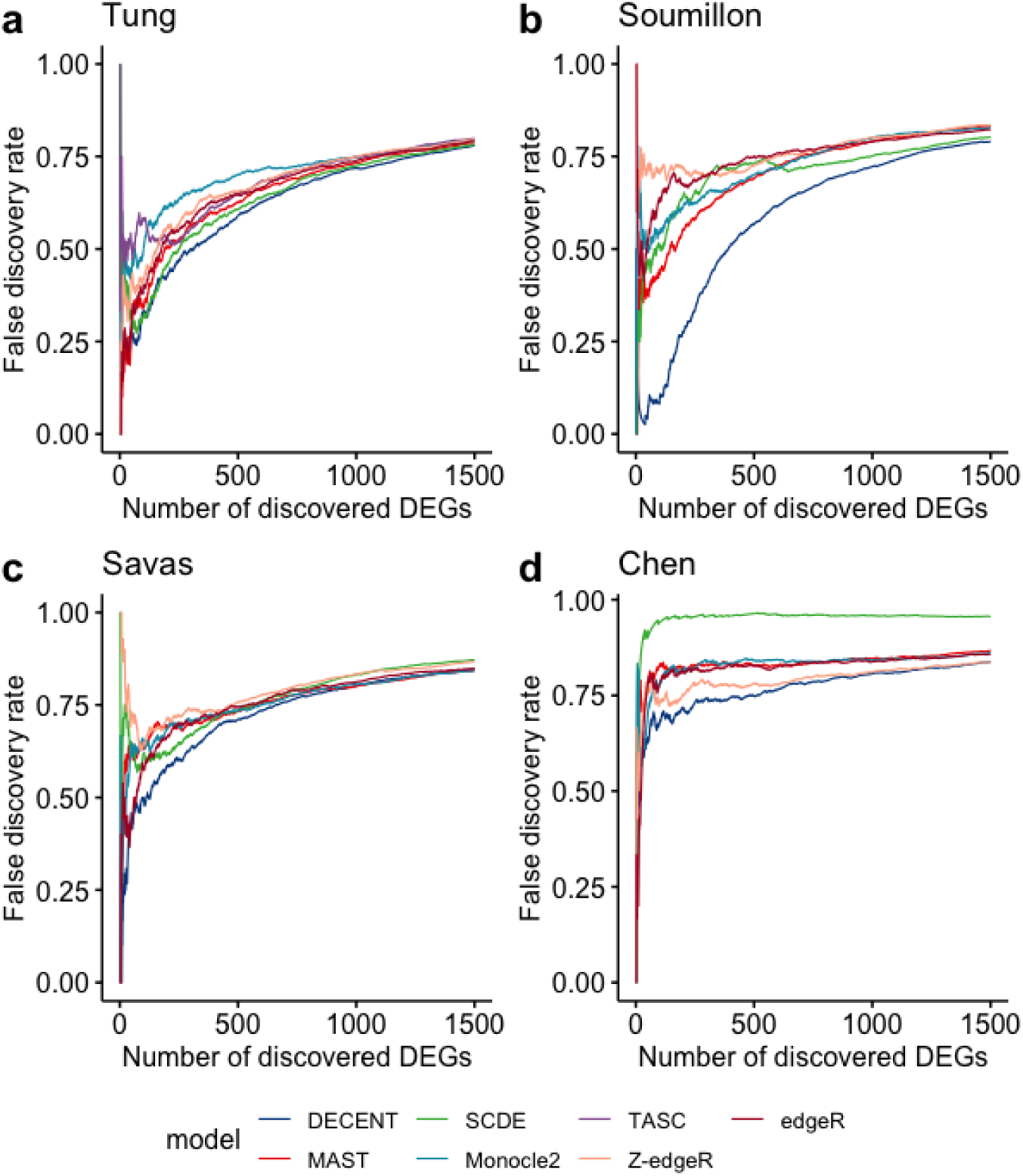
False discovery rate curves for differential expression methods on real datasets. Evaluating the performance different method by false discovery rate (FDR) curves using (**a**) Tung et *al.*, (**b**) Soumillon *et al.*, (**c**) Savas *et al.* and (**d**) Chen *et al.* datasets. Bulk DEGs were considered as conditional positives. DECENT consistently showed the lowest number of false discoveries at the same number of declared DEGs across all four datasets. Again only the top one thousand DEGs were considered to focus on the region of interested. DE methods are denoted by different colors. Z-edgeR denotes ZINB-WaVE-adjusted edgeR. TASC requires spike-ins and was only evaluated using Tung *et al.* data.

ERCC spike-ins were available in Tung *et al.* data. We thus used capture efficiencies estimated from spike-ins for the result shown. This dataset also enabled us to examine how specifying the ranked random capture efficiencies impacts DE performance on real data. We performed DECENT DE analysis again using the ranked random capture efficiencies specifying the range as half, the same and 1.5 times the range of the spike-in estimates. The results turned out to be in concordance with the simulation studies. Although optimal performance was achieved when capture efficiencies estimated from spike-ins were used, there were only small decreases in performance when using the ranked random capture efficiencies (Supplementary Fig.9). This convincingly demonstrated the viability of using the spike-in capture efficiencies for endogenous RNA and that DECENT’s DE performance is also robust to misspecified capture efficiencies.

For the Soumillon *et al.* data, the median of the log fold-change estimates deviates from zero when the standard MLEs were used to estimate the cell size factors *s*_*j*_. This default size factor estimator effectively performs library size normalization on the pre-dropout counts *y*_*i* *j*_. The bias greatly reduced when using the trimmed mean of M values (TMM) method (Robinson and Oshlack, 2010), to estimate the size factors instead and the overall performance of DECENT was slightly improved (Supplementary Fig.10). This suggested that different datasets tend to require different normalization strategies, and suggested the flexibility of our method with regards to normalization strategy.

The benchmarking so far was based on two group comparisons. DECENT performs statistical tests under the under the well-established generalized linear model (GLM) framework and can readily accommodate more complex experimental designs. The Soumillon *et al.* data is a time course experiment, with three time points involved in adipose stem cell differentiation. This allowed us to have a glance at how different DE methods perform on more complex UMI-based scRNA-seq experiments beyond two-group comparisons. We tested the hypothesis that expression of a gene is constant across the three time points. Except for SCDE, which is designed only for two group comparison, and TASC, which requires spike-ins, other methods were compared in this setting. The reference genuine DEGs across the three time points were also derived from the matching bulk experiments. DECENT again outperformed all other methods with an even more pronounced advantage (Supplementary Fig.11).

### Controlling type I errors

Finally, we examined the ability of DECENT to control type I errors. Towards this end, we created a scenario where no genuine DEGs are expected, thus all discovered DEGs are false positives. We randomly split the 221 cells from individual NA19239 in Tung *et al.* data in two groups of sizes 110 and 111. Since the split is random, no biological variation would be expected between the two group of cells, on average. Then the same set of DE methods as above were used to perform DE analysis comparing the two groups. The null hypothesis should hold true for all the genes and hence the nominal p-values obtained from each method should be uniformly distributed. As shown in the quantile-quantile plots, most of the methods produced p-value distributions as desired, including DECENT both with and without using the spike-ins. Only SCDE was producing a conservative p-value distribution with a spike at one, and the p-values from Monocle2 were skewed towards the lower end (Fig.6a). This suggests the negative binomial model fitted directly to the observed data as used in Monocle2 is not able to adjust for the extra variability in the molecule capturing process, thus producing false positives.

**Figure 6:**
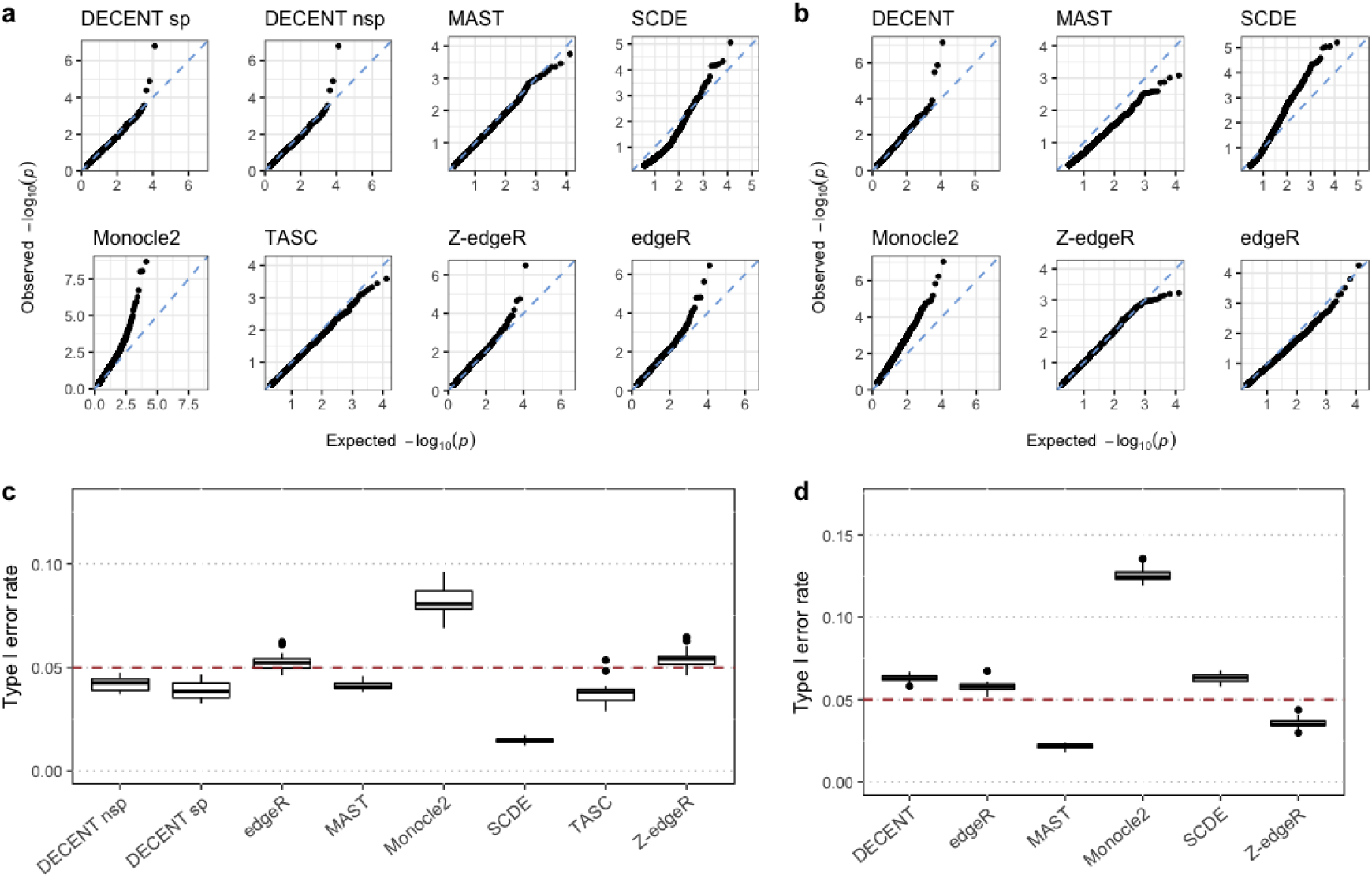
Controlling Type I error rate. We evaluated nominal p-value distributions and type I error rates produce by differential expression methods in absence of genuine DEGs. In panels (**a**) and (**c**), nominal p-values were obtained by comparing two random split group of cells from the NA19239 cell line in the Tung *et al.* dataset. The random split and comparison was performed 20 times. For panels (**b**) and (**d**), nominal p-values were produced by different methods on two randomly sampled groups from stage 3 day 0 cells in the Soumillon *et al.* data. The sampling and comparison was again performed 20 times. (**a**) (**b**) shows quantile-quantile plots of nominal p-values produced by different methods comparing the quantiles of their distribution with the uniform distribution. (**d**) shows observed type I error rates by using a p-value cut-off of 0.05 on nominal p-values produced by different DE methods. Each box was generated based on the same comparisons (n=20) using for both datasets. DECENT nsp denotes DECENT without using spike-ins to estimate capture efficiencies. Overall, DECENT exhibits normal p-value distributions and reasonable control of type I errors in both case.

We conducted the comparison on twenty random splits of the cells. To perform an overall assessment, we calculated the observed proportion of declared DEGs by each method using a p-value cut-off of 0.05. This proportion equals type I error rate and is supposed to match the nominal p-value cut-off on each random splits. The results coincided with that shown in the single split case. Most methods consistently produced observed type I error rates close to 0.05, whereas SCDE was overly conservative and Monocle2 produced the largest number of false positives (Fig.6c).

We also carried out a similar analysis using the Soumillon *et al.* data. In each comparison, we randomly sampled two groups of 200 cells from day 0. And again twenty comparisons were conducted. Given the different features of this dataset, such as being more sparse, MAST exhibited overly low type I error rates with p-value cut-off 0.05, whereas DECENT still showed acceptable control of false positives (Fig.6b, 6d). This could be that the hurdle model used by MAST was overly adjusting for the observed zeros without distinguishing between dropouts and real biological ones. SCDE appeared to have more reasonable observed type I error rate in this case but a closer examination on p-value distribution revealed the same concerns as previously (Fig.6b).

## Discussion

We presented DECENT, a novel statistical method for performing DE analysis on UMI-based scRNA-seq data. The UMI-count data has provided us with a great chance to model the molecule capturing process. The technical variation occurring in this process is precisely characterized by gene and cell-specific beta-binomial dropout models. We were able to perform DE analysis on the inferred pre-dropout data where most technical variation was removed, and hence achieving superior performance. We demonstrated the flexibility of our model for being usable either with or without spike-ins and compatible with different normalization strategies. Also, the model is based on the established GLM theory thus capable of analyzing complex designed experiments. We tested model under the three group one-way ANOVA setting and obtained promising results. Adding more cell-level covariates would also be relatively straightforward (see Methods) and this is catered for in our software.

External RNA spike-ins, such as ERCC spike-ins (Jiang et al., 2011) are a good approach of measuring the technical variation in scRNA-seq data. We use them to estimate capture efficiencies in our model when available. They have also been used in some other scRNA-seq methods (Lun et al., 2017) (Jia et al., 2017). However, given the different features of external RNA spike-in molecules compared with endogenous transcripts such as poly(A) stretch and sequence length, it has been previously found that the amount of technical variation, such as the magnitude of capture efficiencies (Svensson et al., 2017), differs between the two types of molecules. Therefore, models estimated using spike-ins may not be entirely appropriate for endogenous transcripts. How to effectively make use of spike-ins is still a challenging topic in scRNA-seq data analysis. Efforts were made in looking for stably expressed genes in data to substitute for spike-ins (Lin et al., 2017) (Yip et al., 2017). We have used ERCC spike-in mainly as a tool for exploring. If we consider spike-ins as a separate groups of molecules that have a similar capture process to the endogenous RNA molecules, we can then use the same dropout parameters estimated using spike-ins when dealing with endogenous genes. But our method is also flexible enough and allows some of the dropout parameters to differ between the spike-ins and endogenous genes to reflect potential differences in the capture process of the two types of molecules.

In our initial investigation of the dropout model, we found extra variation in the data compared to the cell-specific binomial dropout model. This extra variation is more likely to be spike-in-specific biases rather than random noise (Supplementary Fig.2). However, unlike cell-specific capture efficiencies, the estimated spike-in-specific biases cannot be applied to endogenous genes. Also, we are not able to estimate the gene-specific bias using gene abundance because it is not separable from actual gene mean expression. The separation is only achievable if extra information other than transcript abundance is available. For example, it is plausible that capture efficiencies would depend on gene sequence features such as GC-content and the length of the poly(A) stretch. A more refined dropout model might be built by modeling the relationship between these gene-specific features and the gene-specific biases of capture efficiency.

Although multilevel models with EM algorithm are intrinsically computationally intensive, DECENT has achieved acceptable speed with a series of acceleration approaches such as a gaussian quadrature approximation for large integration and parallelization of all the main steps. For instance, our 500 cells with 3,000 genes simulated data took ∼18 minutes and the largest Chen et al. data with 6,875 cells and 12,929 genes took ∼8 hours to finish on a 28-core XENON Radon Duo R1885 server node with Intel(R) Xeon(R) E5-2690 v4 CPUS @ 2.60GHz.

Some existing models for scRNA-seq allow differential tests beyond the conventional DE analysis, such as testing on differences in the zero fraction, biological variation or even the overall distribution (Korthauer et al., 2016) (Wu et al., 2018) (Wang et al., 2018). But there is still difficulty in assessing the performance such as accuracy, type I error control, etc. of these types of tests due to lack of ground-truth. The smFISH technology is under rapid development. It is able to produce accurate measurements of the biological variation and the zero fraction. As the amount of data and number of genes profilable increases, this should provide us with an opportunity to assess these tests objectively. While DECENT focuses on performing a reliable statistical test for the conventional DE of the mean, it could be extended for performing other types of tests in relatively straightforward manner, given its general modeling framework. For example, we can also model the zero-inflation parameter in the pre-dropout distribution as a function of cellular groups through a logistic linear regression model and test for differences in inflated biological zeros. However, some alteration of the parameter estimation strategy might be needed to achieve valid testing results.

## Methods

### Model formulation

DECENT assumes that unique molecular identifiers (UMI) (Islam et al., 2014) have been used in the scRNA-seq experiment for counting molecules. To permit separation of biological from technical variations, we first assume that in an idealized setting where all molecules are captured, the observed count *y*_*i* *j*_ for gene *i* in cell *j* can be modeled as a zero-inflated negative binomial (ZINB) random variable with parameters θ_*i* *j*_ = (*π*_0*i*_, μ_*i* *j*_, *s*_*j*_, *Ψ*_*i*_), where *π*_0*i*_ is a gene-specific zero-inflation parameter, *Ψ*_*i*_ is a gene-specific dispersion parameter, *μ*_*i* *j*_ is the gene-specific and cellular group-specific mean parameter and *s*_*j*_ represents the size factor for cell *j* that measures differences in the amount of starting material, namely total mRNA between cells.

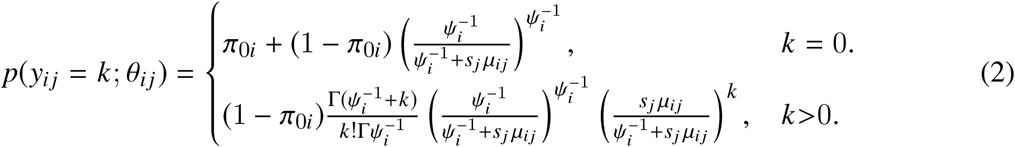

The first line gives the probability of a biological zero. For lowly expressed genes with small mean parameter *μ*_*i* *j*_, the contribution from the second component can be considerable, but for higher abundance genes, the probability of a biological zero largely depends on *π*_0*i*_, with larger values of this parameter being closely associated with higher probabilities of a biological zero.

The gene-wise mean parameter *μ* = (*μ*_*i* *j*_) is assumed to depend on the cell type or group through a log-linear model

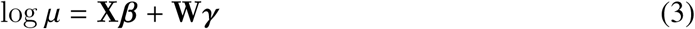

where **X** is the design matrix providing group information and ***β*** are the coefficients. For the completeness of a generalized linear model framework, we also allow including cell-wise covariates **W** to remove unwanted variation (e.g batch effects, cell-cycle phases, etc.). In the most common two group comparisons, we have

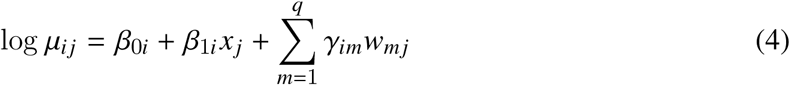

where *x*_*j*_ is simply the binary indicator of cellular group and *β*_1*i*_ has interpretation as the log-fold change (logFC) parameter for gene *i*.

In reality, *y*_*i* *j*_ is unobservable. Instead we have the observed counts *z*_*i* *j*_ that are what remains of the *y*_*i* *j*_ after dropout. DECENT uses a modified beta-binomial distribution to model the capturing process (see Results). We suppose, given *y*_*i* *j*_ as the unobserved pro-dropout molecule count, that the observed count *z*_*i* *j*_ follow a beta-binomial distribution

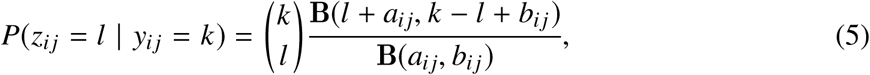

where **B**(., .) is the Beta function. We reparametrize the model by

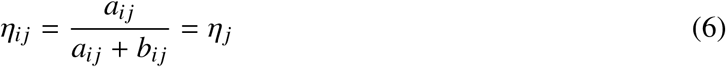

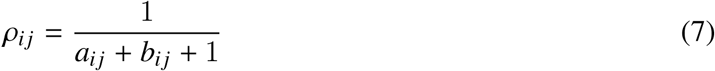

where we suppose *η*_*i* *j*_ does not depend on *i*, and so *η*_*j*_ represent the cell-specific capture efficiency in cell *j*. The amount of variability within the cell is measured by the dispersion parameter *ρ*_*i* *j*_ that depends on the mean expression of gene *i* in cell *j* via a cell-specific linear model:

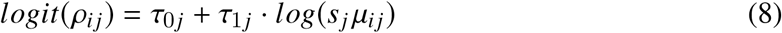

Capture efficiencies *η*_*j*_ are estimated using spike-ins when available. We also provide a strategy to produce functional capture efficiencies when spike-ins are not available. These will be discussed in the following sections. Since the pre-dropout counts for endogenous genes are unobserved, we use an Expectation-Maximization algorithm to estimate the gene-specific parameters θ_*i*_ = {*π*_0*i*_, *β*_0*i*_, *β*_1*i*_, <u>γ</u>*i*, *Ψ*_*i*_ } and cell-specific parameters *s*_*j*_. Not surprisingly, the E-step involves evaluating the conditional probability of an observed zero count being a biological zero, 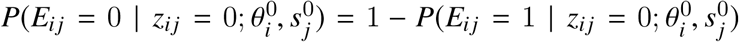, where *E*_*i* *j*_ is a binary indicator of gene *i* being truly expressed in cell *j*, i.e. *y*_*ij*_ > 0 (see Supplementary methods for details). The **τ _*j*_** = (τ_0*j*_, τ_1*j*_) parameters in the dropout model are estimated during the EM iterations using either spike-ins or endogenous gene counts. If endogenous genes are used to estimate **τ _*j*_** it would allow some dropout parameters to be different between the spike-ins and endogenous genes, reflecting inherent differences in their dropout processes.

### Estimating capture efficiencies

Spike-in data are used to estimate the capture efficiencies when available. Suppose we added *n* spike-ins at the known concentrations *c*_1_, *c*_2_, … *c*_*n*_ into cell *j* and subsequently observe *z*_1*j*_, *z*_2*j*_, … *z*_*nj*_ molecules respectively. The cell-specific capture efficiency for any cell *j* is estimated as the proportion of molecules observed after sequencing relative to the total number of molecules initially added:

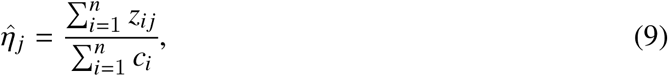

This is the method of moments estimator (MME) of *η*_*j*_ under the beta-binomial-Poisson model for spike-ins (see Supplementary Methods).

Many scRNA-seq data do not have spike-ins. Also, although the spike-in capture efficiencies can be used as a good approximation, they may not be exactly the same as those of endogenous RNA. Interestingly, we found that if we specified a set of inexact capture efficiencies, other components of the model will compensate for the inaccuracy and produce DE results almost as reliable as if we had the correct values. This is due to a property of the our model that is explained below: +

Let *Y* be the pre-dropout count where *Y* ∼ ZINB(*π*_0_, *s μ*, *Ψ*). Given *Y*, the observed data *Z* follows Beta-Binomial distribution, *Z* | *Y* = *y* ∼ *BB*(*y*, *a*, *b*). This is the usual parametrization of beta distribution where 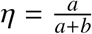 and *ρ* = (*a* + *b* + 1)^-1^. It turns out that the marginal distribution *F*_*Z*_ of *Z* can be approximated by the marginal distribution *F*_*Z*_*′* of *Z*^*′*^, i.e.

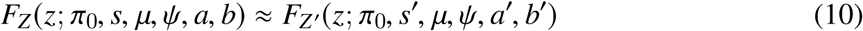

where *Z*^*′*^ | *Y* ^*′*^ = *y*^*′*^ ∼ ***BB***(*y*^*′*^, *a*^*′*^, *b*^*′*^) and *Y* ^*′*^ ∼ ZINB(*π*_0_, *s*^*′*^μ, *Ψ*). When we misspecify capture efficiency *η* as *η*^*′*^, then *a*, *b*, and *s* correspondingly become 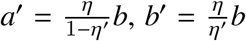 and 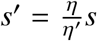 to keep a similar marginal distribution. This is illustrated in Supplementary Fig.6.

The above result means that if we misspecify the capture efficiency by using *η*^*′*^ rather than *η*, the misspecification can be approximately corrected by scaling the size factor estimates accordingly. The remaining effect will be compensated by adaptive estimation of τ. Certainly it is still preferable to get capture-efficiency estimates as close as possible to the true value. Motivated by the above results and our experience with real datasets showing that capture-efficiency is the biggest factor contributing to the variation in the observed library sizes, we devised a method for generating functional capture efficiencies when spike-ins are not available:

This method requires the range of capture efficiency be supplied. Let the lower and upper bounds of this range be min_*η*_ and max_*η*_, respectively. The cell-specific capture efficiencies are specified as follows:

- Compute library size for each cell and denote the log10 of these by *L*_1_, *L*_2_, … *L*_*N*_. To minimize the impact of a few genes having very large counts, we can also use trimmed sums instead of full sums here. Denote the minimum and maximum log10 library size as *L*_*min*_ and *L*_*max*_.
- Calculate weight for cell *j* as 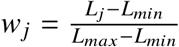.
- Estimate the capture efficiency for cell *j* as (1 - *w*_*j*_) min_*η*_ +*w*_*j*_ max_*η*_. This ensures that cells with larger library size will have larger capture efficiency and the capture efficiency estimates are bounded within (min_*η*_, max_*η*_) interval.

We refer to this as the ranked random capture efficiency.

## Estimating the parameters τ _*j*_

Besides the capture efficiencies, the parameters **τ _*j*_** in the logistic model for the beta-binomial dispersion parameter are also crucial for the dropout model. Like capture efficiencies, we can opt to use **τ _*j*_** estimated from spike-ins for endogenous genes, when spike-ins are available. But we can only estimate **τ _*j*_** using endogenous gene counts when spike-ins are not available. We found that the **τ _*j*_** estimates for endogenous genes often differ from those for the spike-ins in real scRNA-seq data. Therefore, we strongly advise users to estimate the parameters **τ _*j*_** using endogenous gene counts. This also make the model more robust to misspecification of capture efficiencies, as the 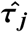 will now account for the variation due to inaccurate capture efficiencies. However, estimating cell-specific 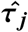 using endogenous genes can be a difficult task especially for sparse data with low counts or large zero fractions. We therefore implemented two options for obtaining the 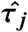 estimates. The first one is to assume 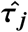 are constant across cells, resulting two global parameters (τ_0_, τ_1_). Under this assumption, we have

Within each EM iteration, after the M-step (see Supplementary methods)

Given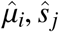, *ŝ*_*j*_ and capture efficiency estimates 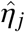, the correlation parameter for each gene *i* is estimated by maximizing

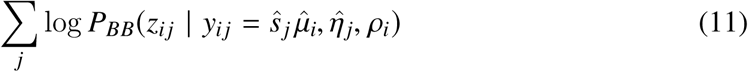

where ***P*_*BB*_** is the Beta-Binomial density with probability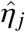, size parameter 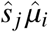 and dispersion parameter *ρ*_*i*_.

The τ_0_ and τ_1_ estimates are updated as the intercept and slope estimates of the following regression model:

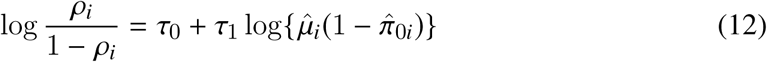

When we have enough information in the data, the other option is to estimate cell-specific 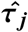 by

Given 𝔼 (*y*_*i* *j*_ | *z*_*i* *j*_) from the E-step and CE estimates 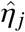, for each cell *j*, the cell-specific parameters τ_0*j*_ and τ_1*j*_ are updated by maximizing the following log-likelihood

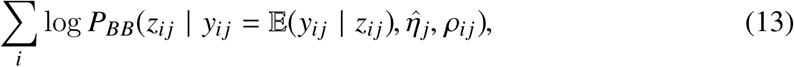

where *ρ*_*i* *j*_ is a function of τ_0*j*_ and τ_1*j*_ through

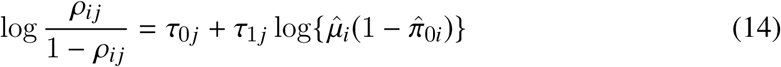

### DE analyses

Differential expression across two cellular groups for the *i*^*th*^ gene is assessed by testing the hypotheses:

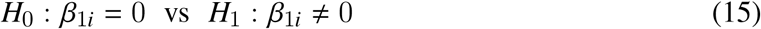

using the likelihood ratio test (LRT) statistic,

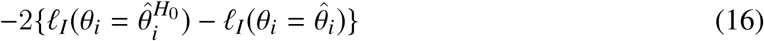

where 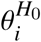 is the maximum likelihood estimator (MLE) of θ_*i*_ under the restriction that 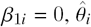 is the MLE under the unrestricted model and *ℓ*_*I*_ is the log-likelihood of the observed incomplete data *z*_*i* *j*_. For simple two cell-type comparisons, the statistic is approximately distributed as 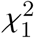 under *H*_0_. More generally, when performing DE across *p* different cell-types or conditions, the statistic is approximately distributed as 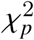 under *H*_0_.

### Public datasets

- **Tung *et al.* dataset**: This dataset is from the (Tung et al., 2017) benchmarking scRNA-seq experiment. We downloaded the filtered UMI count matrix from their GitHub repository (https://github.com/jdblischak/singleCellSeq). The full dataset contains three Yoruba (YRI) induced pluripotent stem cell (iPSC) lines, with three 96-well plates per individual. ERCC spike-ins (Jiang et al., 2011) and UMI were used. Each replicate was also used to generate a matching bulk RNA-seq sample. We only used data of two individuals NA19101 (201 cells) and NA19239 (221 cells) for the analyses. Reference genuine DEGs were derived by selecting the 500 DEGs with smallest p-values produced by *limma-voom* (Ritchie et al., 2015) *using the bulk RNA-seq samples of the two individuals.*
- ***Soumillon et al.* dataset**: This dataset is publicly available from Gene Expression Omnibus (GEO) repository GSE53638. Cells were collected at different stages and different time points of directed differentiation of human adipose-derived stem/stromal cells (Soumillon et al., 2014). FACS sorted cells were sequenced using the SCRB-seq protocol with UMI. To benchmark DE methods using a two group comparison, we compare the stage-3 differentiated cells at day 0 (baseline, 943 cells) versus day 7 (1006 cells). All three time points day 0, day 3 (1019 cells) and day 7 of stage-3 differentiated cells were used for the three group DE analysis. Cells have been filtered by the authors and genes with log total UMI counts over one median absolute deviation (MAD) lower than the median were removed after subsetting the cells. The matching bulk RNA-seq data have only one sample per time point. Therefore, we selected the 500 genes with the largest log fold-change as the reference genuine DEGs for two group comparison. We also used the 500 genes with largest variances across three time points for benchmarking the three group analysis. Log fold-changes and variances were calculated based on log count per million (CPM) with high prior count 5 for stabilization.
- **Savas *et al.* dataset**: The experiment profiled the transcriptomes of tumour infiltrating T cells from triple-negative breast cancer patients. The full dataset is available from GSE110686. Pre-processing and cluster analysis were performed as described in (Savas et al., 2018). We used the CD8^+^ TRM (606 cells) and CD8^+^ non-TRM (1097 cells) clusters (CD8^+^γd together with CD8^+^ effector memory) as the two groups to be compared. Data from case one was used in the analysis. A corresponding bulk RNA-seq experiment is available from GSE110938, comparing CD8^+^CD103^+^ and CD8^+^CD103^-^ FACS sorted populations. The bulk DEGs used as our reference gene list is available as a supplementary table in the original paper.
- **Chen *et al.* dataset**: The single-cell and bulk RNA-seq data are both available from GEO entry GSE113660. The scRNA-seq experiment profiled over six thousand cells from the Rh41 cell line. After quality control and cluster analysis, two clusters representing respectively CD44^+^ (3074 cells) and CD44^-^ (3801 cells) populations were obtained. The cluster labels were acquired through personal contact with the authors. Genes with total UMI count less than 100 were filtered out. The matching bulk RNA-seq data has three batches. Each batch contains a CD44 high, a CD44 low and an unsorted sample obtained via FACS-sorting. The top 500 DEGs comparing CD44 high and CD44 low samples were used as reference DEGs.
- **Zeisel *et al.* dataset**: The experiment sequenced three thousand cells in the mouse somatosensory cortex and hippocampal CA1 region. Cells were classified into two levels of cell types. The dataset is available from the authors via: http://linnarssonlab.org/cortex. We only used the cells within the “pyramidal CA1” level 1 class for our analysis. This dataset exhibits very high proportions of spike-in counts in most cells. This suggests intense competition between the spike-in and endogenous molecules for read counts and the quantification of endogenous genes is likely to be affected. To moderate this, we removed cells with more than 50% UMI counts coming from spike-ins when fitting the model for endogenous genes (remaining 932 cells). We further filtered out genes with total UMI counts over one MAD lower than the median.
- **ERCC spike-in datasets**: Apart from the ERCC spike-in data within the Tung *et al.* and Zeisel *et al.* dataset, three other datasets were downloaded from their NCBI GEO repositories: GSE54695 (Grun *et al.*), GSE65525 (Klein *et al.*) and GSE63473 (Macosko *et al.*). The Zheng *et al.* ERCC spike-in experiment is available on the 10x Genomic website: https://support.10xgenomics.com/single-cell-gene-expression/datasets/1.1.0/ercc. All spike-in datasets underwent the same filtering steps. We removed spike-ins that have nominal count < 0.05 or a mean observed count higher than the nominal count. We filtered out cells with total UMI counts more than 2 MADs below the median total UMI count.
- **osmFISH dataset**: The authors applied a newly developed cyclic single-molecule FISH protocol, termed ouroboros smFISH (osmFISH) to cells from the mouse somatosensory cortex tissue. The experiment quantified RNA molecules from 33 target genes in more than four thousand cells in a brain tissue section. The smFISH count data and cluster labels are available at http://linnarssonlab.org/osmFISH/.

### Fitting the dropout models using ERCC spike-ins

We use a Poisson distribution to model the pre-dropout molecule count of spike-ins:

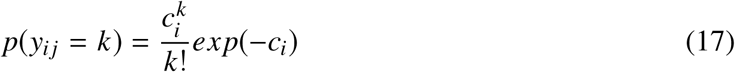

Under both the binomial and beta-binomial dropout models, the capture efficiency *η*_*j*_ is estimated as in (10) above. It is either the MLE or MME for *η*_*j*_ (see Supplementary Methods). In our investigation of the beta-binomial dropout model, we also needed to estimate the parameters *ρ*_*i*_ or **τ _*j*_** in each cell-wise model. To take into account this Poisson variation in the estimation of **τ _*j*_** for spike-ins, we first simulate the unobserved count 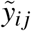 under Poisson(*c*_*i*_). Each spike-in was simulated 50 times to achieve stable estimation. Then the the MLEs 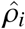 or 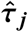 under either 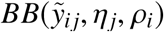 or 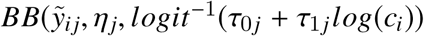 can be obtained by maximizing the beta-binomial likelihood function.

### Calculating the deviances for the binomial and beta-binomial dropout models

Under the binomial dropout model, the distribution *z*_*i* *j*_ is another Poisson distribution with rate *η*_*j*_ *c*_*i*_ because the binomial thinning of Poisson is still Poisson (Casella and Berger, 2002). Therefore, the binomial deviance for spike-in *i* in cell *j* is simply:

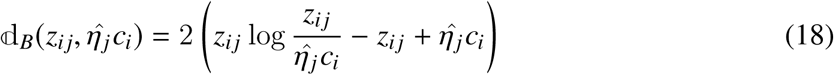

Under the beta-binomial dropout model *z*_*i* *j*_ | *y*_*i* *j*_ ∼ *BB*(*η*_*j*_, *y*_*i* *j*_, *ρ*_*i* *j*_), the deviance for spike-in *i* in cell *j* is given by,

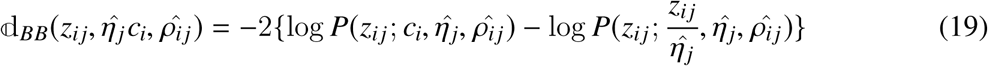

where *P*(*z*_*i* *j*_; *c*, *η*, *ρ*) =Σ*y P*(*z*_*i* *j*_ | *y*_*i* *j*_; *η*, *ρ*)*P*(*y*_*i* *j*_; *c*) is the marginal probability distribution of the observed data. Here *P*(*z*_*i* *j*_ | *y*_*i* *j*_; *η*, *ρ*) and *P*(*y*_*i* *j*_; *c*) are the beta-binomial and Poisson probability mass function (PMF). In practice, the marginal distribution was calculated numerically using Gaussian quadrature that approximates the summation as integration with a continuity correction. Then the deviances for cell *j* models are

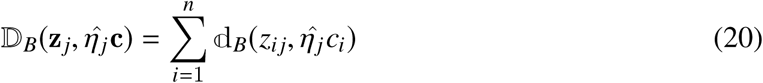

Or

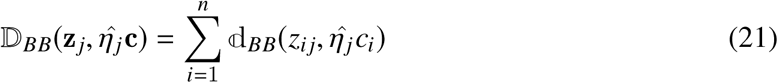

which asymptotically follow 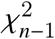 and 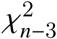, respectively, under the null hypothesis.

### Testing for overdispersion and zero-inflation in the smFISH data

We denote the smFISH molecule count of gene *i* in cell *j* by *y*_*i* *j*_, as it is supposed to be a accurate quantification of the actual RNA count without dropout. To investigate the pre-dropout distribution, we fitted three models: *Poisson*(*s*_*j*_ μ_*i*_), ***NB***(*s*_*j*_ μ_*i*_, *Ψ*_*i*_) and ***ZINB***(*s*_*j*_ μ_*i*_, *Ψ*_*i*_, *π*_*i*_) to the (*y*_*i* *j*_), where *s*_*j*_ is the cell-wise size factor with the restriction 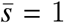, *μ*_*i*_ is the gene-wise mean parameter, *Ψ*_*i*_ is the gene-specific NB dispersion parameter and *π*_*i*_ is the gene-specific zero-inflation parameter. Under all three models, the parameters *s*_*j*_ can all be estimated by MLE 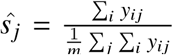, where *m* is the number of cells. This allowed us to fit gene-wise models easily using the R GLM framework with the 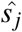 supplied as offsets. We used the *glm* function from the stats package to fit the Poisson models, the *glm.nb* function from the MASS (v7.3-50) package for fitting the NB models and the *zeroinfl* function in the pscl package (v1.5.2) for the ZINB models. To test for overdispersion in each gene, we used the Cameron and Trivedi’s score test (Cameron and Trivedi, 1990) on the fitted gene-wise Poisson model. We used the *dispersiontest* function implemented in the AER(v1.2-5) R package with NB2 as the alternative model. As for testing zero-inflation, we performed a likelihood-ratio test between the fitted NB and ZINB model of each gene. Note that the null distribution in this case is asymptotically 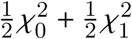 rather than 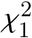 since the null hypothesis *π*_*i*_ = 0 is on the boundary of the parameter space [0,1].

## Simulation

We simulated data for 500 cells belonging to two different cell types (224 vs 276 cells for each type). For each cell, the observed count data for 3000 endogenous genes and 92 ERCC spike-ins are generated from a zero-inflated negative binomial (ZINB) model for the pre-dropout count. For each gene, the gene-specific mean and dispersion parameters are sampled randomly from the empirical distribution of these parameters in Tung’s data for NA19101 and NA19239 cell lines. Because Tung’s data contains atypically low percentage of zero counts for scRNA-seq data (∼35%), the mean parameter for our simulation studies is scaled by a factor of 0.1, resulting in approximately 80% zero counts in the dataset. Approximately 10% of the genes are designated as DE genes and their fold-change parameters are randomly generated from Gamma(2,2) distribution. For non DE gene, the fold change parameters are set to 1.

Biological zeroes are added through zero-inflated parameter *π*_0_, generated from Beta (3,17) distribution, which results in an average of 15% biological zeroes in the pre-dropout counts. The capture efficiency (CE) parameters are also generated from the empirical distribution of CE in data for NA19101 and NA19239 cell lines. Once the pre-dropout counts are simulated, the observed counts are generated by applying Beta-Binomial dropout model to the pre-dropout counts. Global dropout parameters are used with τ_0_ = −1.5 and τ_1_ = −0.3. Finally, the size factor parameters are generated separately for the two cell-types so on average the first cell type has smaller size factor than the second. This is achieved by generating the size factors for the first cell type from (scaled) Gamma (4,5) distribution and for the second cell type from (scaled) Gamma (5,4) distribution. The scaling factors are chosen so that the average size factors across all cells is equal to 1. Before performing the benchmarking, we removed low abundance genes that are expressed in less than 3 cells.

## Performance evaluation

The performance of different methods for identifying genuine DEGs was evaluated using the partial Receiver Operating Characteristic (pROC) curve of true positive rate (TPR) plotted against false positive rate (FPR) within the range of FPR < 0.1 and false discovery rate (FDR) curve showing the FDR among the top n discovered DEGs. These rates are defined as:

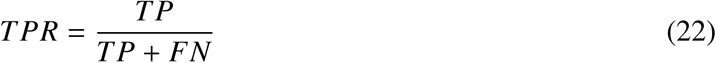

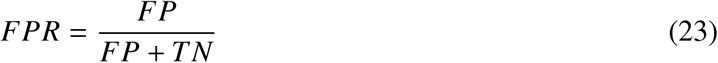

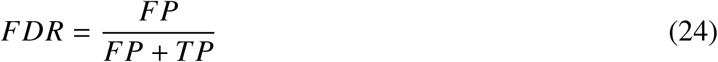

where TP, FP, TN and FN denote number of true positives, false positives, true negatives and false negatives, respectively.

## Benchmark settings

DECENT were run with its default parameters on the Soumillon *et al.* and Savas *et al.* datasets. Cell-specific estimation of **τ _*j*_** was used in *Tung et al.* data and disabled in all the other data. This is because an acceptable amount of information in the data is required in order to obtain reliable cell-specific **τ _*j*_** estimates. Generally we suggest trying cell-specific estimation only on datasets having less than ∼70% observed zeros, and ideally with spike-ins to estimate the capture efficiencies. We increased the range of ranked random capture efficiencies for Chen *et al.* data from the default [0.02, 0.1] to [0.04, 0.2] given its high counts. We used the default settings for MAST (v1.6.1), Monocle2 (v2.8.0), TASC and edgeR (v3.22.2). Log CPM with prior count 1 was used as input for MAST and the likelihood ratio test is used in edgeR. As for SCDE (v1.99.2), we set *min.count.threshold* to 1, increased *min.nonfailed* to 10 as suggested by the authors for using it on large-scale UMI data. For ZINB-WaVE (v1.2.0), the performance appears to be very sensitive to the parameter epsilon, and so we selected the optimal epsilon parameter for each dataset from a range of 10^3^ to 10^13^. The groups to be compared were supplied as cell-level covariate X. Other parameters including those in the following weighted edgeR analysis were left as default.

To derive reference DEGs, we used default settings of limma-voom (v3.36.1) for the DE analyses of the Tung *et al.* and Chen *et al.* matching bulk RNA-seq data. In these two cases, we retained genes with cpm > 1 in more than 3 samples. For the Chen *et al.* bulk data, a batch dummy variable was included in the design matrix to perform paired comparisons. For the Soumillon *et al.* bulk data, we retained genes with non-zero measurements in both day 0 and day 7 samples for two group comparison, and those which are positive in all three time points for the three group comparison. The top DEGs were inferred by ranking in terms of log fold-changes or variances as describe above.

The R scripts used for the analyses are available via GitHub: https://github.com/cz-ye/DECENT-analysis.

## Software availability

DECENT is implemented as a R package and available from the GitHub repository: https://github.com/cz-ye/DECENT.

## Author Contributions

A.S. and T.P.S conceived the idea and developed the methods. A.S. and C.Y. designed and developed the software. A.S. and C.Y conducted the simulation studies and C.Y conducted the real data analyses. All authors contributed to interpretation of results, writing the manuscripts and approved the final submitted version of the manuscript.

